# Cancer-on-a-chip model shows that the adenomatous polyposis coli mutation impairs T cell engagement and killing of cancer spheroids

**DOI:** 10.1101/2023.10.17.562521

**Authors:** Valentin Bonnet, Erik Maikranz, Marianne Madec, Nadia Vertti-Quintero, Céline Cuche, Marta Mastrogiovanni, Andrés Alcover, Vincenzo Di Bartolo, Charles N. Baroud

## Abstract

Evaluating the ability of cytotoxic T lymphocytes (CTLs) to eliminate tumor cells is crucial, for instance to predict the efficiency of cell therapy in personalized medicine. However, the destruction of a tumor by CTLs involves CTL migration in the extra-tumoral environment, accumulation on the tumor, antigen recognition, and cooperation in killing the cancer cells. Therefore, identifying the limiting steps in this complex process requires spatio-temporal measurements of different cellular events over long periods. Here, we use a cancer-on-a-chip platform to evaluate the impact of adenomatous polyposis coli (APC) mutation on CTL migration and cytotoxicity against 3D tumor spheroids. The APC mutated CTLs are found to have a reduced ability to destroy tumor spheroids compared with control cells, even though APC mutants migrate in the extra-tumoral space and accumulate on the spheroids as efficiently as control cells. Once in contact with the tumor however, mutated CTLs display reduced engagement with the cancer cells, as measured by a new metric that distinguishes different modes of CTL migration. Realigning the CTL trajectories around localized killing cascades reveals that all CTLs transition to high engagement in the two hours preceding the cascades, which confirms that the low engagement is the cause of reduced cytotoxicity. Beyond the study of APC mutations, this platform offers a robust way to compare cytotoxic cell efficiency of even closely related cell types, by relying on a multiscale cytometry approach to disentangle complex interactions and to identify the steps that limit the tumor destruction.

## INTRODUCTION

The adenomatous polyposis coli (APC) gene encodes a large multi-domain protein involved in multiple cellular functions, such as adhesion, proliferation, and differentiation [1–4]. APC is one of the key components of the Wnt/*β*-catenin pathway. Dysregulation of this pathway by sporadic mutations of APC is involved in the emergence of around 80% of colorectal cancers [3]. APC is also mutated in an inherited syndrome called familial adenomatous polyposis (FAP), a disease that leads to the development of hundreds to thousands of intestinal polyps [1]. As these individuals have a mutated allele of APC in every cell, other cell functions are modified. For instance, APC interacts with the cytoskeletal components [5] as it is implicated in actin polymerization and microtubule organization [6, 7]. Therefore the potential impact of APC mutations on immune functions in FAP patients have recently started to be investigated, for instance using an APC^Min*/*+^ mouse model. These mice bear a heterozygous mutation in the murine homolog of the APC gene and have been largely used as a relevant pre-clinical model for this pathology [2, 8–10].

Specifically, perturbations of APC have been shown to affect T cell functions in several ways. First the production of cytokines by regulatory T cells is altered, which impairs their anti-inflammatory functions [11]. Additionally, destabilization of the immunological synapse and reduced delivery of cytotoxic granules have both been observed in APC silenced T cells, resulting in an impaired killing efficiency of cytotoxic T lymphocytes (CTLs) during *in vitro* killing assays [12]. Finally, anomalous migration of T cells from FAP patients was observed during *in vitro* assays. This defect was in part due to impaired integrin-mediated cell adhesion, possibly leading to decreased immune surveillance efficiency and effector function [13, 14]. It is unclear however how these modifications of the immune functions combine together and if the combined effect may reduce their ability to control polyp growth and transformation into cancerous tissue.

This question can be addressed by comparing the ability of APC mutated T cells vs. control cells to reduce tumor size in a model of immune-cancer interactions. A wide variety of killing assays are available, ranging from the killing of two-dimensional monolayers of cells [15, 16] to three dimensional (3D) cancer spheroids or organoids [17, 18] or even animal models [19–21] However, none of the approaches above has the spatial or temporal resolution to observe how modifications in the individual functions (motility, signaling, antigen recognition, killing) combine to yield the observed final behavior, in order to identify which factor, if any, effectively inhibits tumor control.

In this context, the emergence of microphysiological systems (MPS), such as organs-on-a-chip and organoids, provides new tools to study the spatio-temporal dynamics of biological processes. Several different MPS approaches have been developed for cancer applications, particularly for the study of cancer-immune system interactions, as reviewed in several recent articles [22–26]. Recent work has been focused on demonstrating the relevance of MPS models for tumor behavior *in vivo* [27, 28], while engineering advances have involved coupling these models with imaging and image analysis methods to obtain information on the biological and biophysical processes [29–31]. However, none of the existing work has yet provided new mechanistic insights on CTL-cancer interactions in the context of specific diseases.

Here, we build on recent cancer-on-a-chip developments to identify the functional differences between APC-mutated and non-mutated CTLs. This is performed by integrating an experimental and analytical pipeline to measure the destruction of a tumor by the CTLs, using the murine model of B16 melanoma cells expressing Ovalbumin peptide (B16-OVA). The B16 cells were challenged by CD8^+^ CTLs bearing the OVA-specific OT-I transgenic T cell receptor (TCR) [16, 18, 32]. The impact of the APC mutation on the CTL activity was investigated by comparing control CTLs from *APC*^*+/*+^ OT-I mice with CTLs from APC^Min*/*+^ OT-I mice. The immune challenge was performed using a droplet microfluidic platform [31, 33] that enabled the culture of cancer spheroids in a primary hydrogel droplet, followed by the addition of the CTLs at a well defined time-point. While end-point imaging revealed the reduced ability of the APC^Min*/*+^ cells to destroy the tumor spheroids, single-cell tracking allowed us to compare the cell behaviors at different phases of the process. We show that the APC^Min*/*+^ mutation does not reduce the ability of the CTLs to reach the tumor and accumulate on it but instead impairs their ability to engage with the cancer cells.

## RESULTS

### Microfluidic droplets provide a suitable environment to study tumor spheroids and cytotoxic T lymphocytes interactions

The antagonistic interactions between CTLs (APC^+*/*+^ or APC^Min*/*+^) with cancer spheroids were studied in droplets that were anchored within a microfluidic device [34]. This approach has been shown to be suitable for different cell cultures [26, 35, 36], while the use of asymmetric anchor shapes provided a method to bring different cell types into contact at well-controlled times [31, 33]. In the current study, it was important to produce large tumor aggregates, compared to CTL diameter of about 15-20 µm, to ensures sufficient spatial resolution to follow CTL migration on the spheroids. Therefore, a new microfluidic design was implemented in which 0.7 µl droplets were maintained in a chamber containing 68 independent anchors (Fig. 1a and SI Fig. S1a). The chips were fabricated using a 3D printed mold (see SI Fig. S1a), which allowed complex 3D geometries to be molded into a PDMS device (see Materials and Methods).

**Figure 1.**
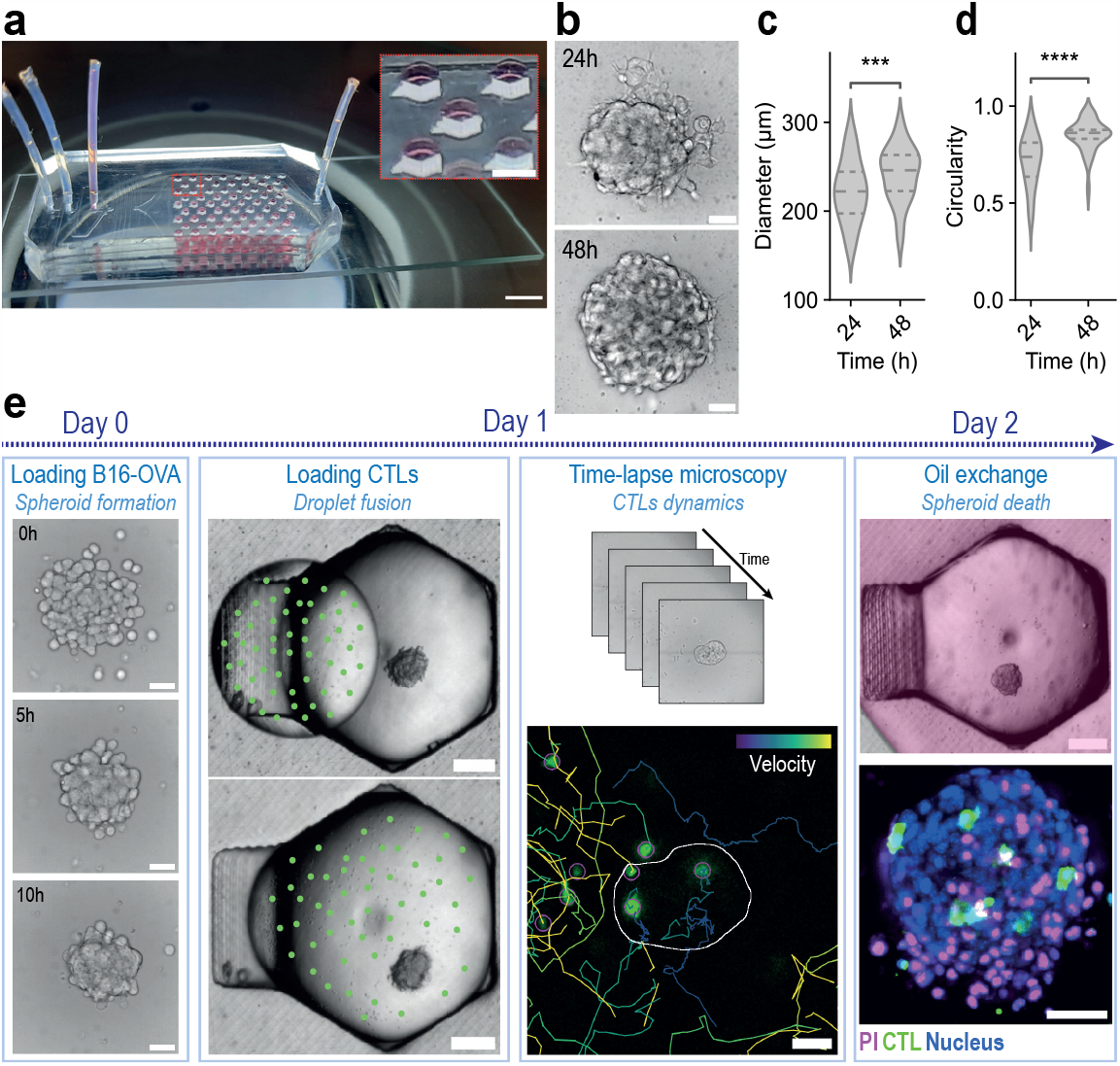
Microfluidic immuno-oncology model and experimental protocol. **(a)** Photograph of the microfluidic chip containing colored water droplets. The PDMS chip is bonded on a standard glass slide. Scale bar 5 mm - Inset: zoom on the droplet anchors. Scale bar 1 mm. **(b)** Representative images of a spheroid in brightfield microscopy at 24 h and 48 h after cell loading. Scale bar 50 µm. **(c)** Distribution of spheroid diameters at 24 h and 48 h. The diameter is computed as 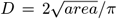. **(d)** Spheroid circularity at 24 h and 48 h. Circularity = 4*π×*area/perimeter^2^. A circularity value of 1.0 indicates a perfect circle. **(c) – (d)** Statistical differences were calculated by two-sided Mann-Whitney-Wilcoxon test (n=90 spheroids) **(e)** Experimental protocol and representative images of the different steps and readouts. Columns from left ro right: (1)B16-OVA cells are loaded in droplets containing culture medium and Matrigel and form a spheroid by aggregation in the first 24 h. Scale bars 50 µm. (2) A second droplet that contains CTLs (represented by green dots) is merged with the first one containing the spheroid. Scale bars 200 µm. (3) Time lapse microscopy is performed and CTL trajectories are extracted in the gel and on the spheroid. Scale bars 50 µm. (4) Oil phase is replaced by an aqueous medium with PI to stain dead cells. Scale bar 200 µm. After incubation, spheroid death can be measured by confocal microscopy. Scale bar 50 µm.

The primary droplets contained around 400 cancer cells each in a mixture of culture medium and Matrigel. The cells formed compact and round tumor aggregates of radii ranging from 150 to 300 µm, when observed 24 h after the beginning of the experiment (Fig. 1b, c). As cell culture in such small volumes can affect cell behavior [26], the viability and proliferation of the cancer spheroids was first characterized without adding CTLs. During 48 h of culture after droplet loading, the spheroids grew due to cell division (Fig. 1b-c) and became more circular (Fig. 1d) due to cell compaction and organization in the aggregate. Cell proliferation within the spheroids was measured after 24 h of culture in the droplet using Bromodeoxyuridine (BrdU) staining (SI Fig. S1b-c). On average 20 % of cells were dividing within a 5 h window, indicating a good proliferating state of the spheroid, since it has been reported that the B16 cells doubling time is around 20 h [37]. Similarly, spheroid survival at 24 h and 48 h was assessed using propidium iodide (PI) and caspase 3/7 dye, showing below 6 % dead cells and below 4% apoptotic cells (SI Fig. S1d-g). Taken together these measurements show that the B16 spheroids could be maintained in a healthy state for at least 48 hours in the droplets without significantly affecting cell viability or proliferation.

The anchor geometry included a tapered protrusion on one side that created a weaker secondary anchoring region. This was then used to trap a smaller second droplet (volume 0.25 µl) containing the CTLs that was added 24 h after initial loading of tumor cells. The sloped ceiling of the secondary anchor ensured that the two droplets were in close contact, allowing them to merge when the surfactant was washed out from the oil. As a result, the CTLs were inserted into the Matrigel droplet containing the cancer spheroid, where their motion and the resulting effect on the spheroid was followed by time-lapse microscopy. At the end of the experiment, the oil was replaced by an aqueous solution containing PI, in order to observe the dead cells after 24 h of coculture. The complete experimental protocol is illustrated in Fig. 1e and key steps can be visualized in Movie S1.

### APC mutation reduces CTL killing efficiency

The impact of the APC mutation on CTL cytotoxic activity was assessed by measuring the reduction in tumor size during coculture of CTLs with tumor spheroids (Fig. 2a), as this feature is often considered when comparing efficiency of different therapeutic strategies against cancer *in vitro* and *in vivo* [32, 38, 39]. Brightfield images to measure spheroid reduction is not informative due to the impossibility of differentiating dead and live cells. However, it is known that during cell apoptosis, actin cytoskeleton undergoes multiple rearrangements correlating with changes in cell shape [40, 41]. Therefore, by staining the actin fibers, it is possible to follow the dramatic reorganization of actin in space and time (Fig. 2a, SI Fig. S2a (white arrow) and Movie S2), and to use it as a marker of the progressive destruction of the cancer spheroid. The validity of this imaging marker was verified by staining the tumor spheroid with PI 24 h after CTLs insertion and confirmed that the region of disorganized actin fibers corresponds to a region of high cell mortality (SI Fig. S2b).

Figure 2b shows the portion of spheroid still alive 5 h and 10 h after the first contact of a CTL with the spheroid, for APC^+*/*+^ (refered from now on as control cells) and APC^Min*/*+^ cells. A significant reduction in CTL killing efficiency is measured in the case of the APC mutant compared with the control. These measurements offer a first indication of a detrimental impact of APC mutation on CTL killing efficiency. Importantly, this difference does not come from a bias in CTL accumulation on the spheroids as shown on SI Fig. S2c. In addition, the percentage of cancer cell death after 24 h of coculture was measured using PI staining and 3D confocal microscopy (Fig. 2c), independently of the actin measurements. The quantification confirms a decrease in cancer cell death under the action of APC^Min*/*+^ CTLs compared to the control (Fig. 2d). Again, the difference in destruction of tumor cells does not come from a bias in CTL number on the spheroid after 24 h of coculture (SI Fig. S1d).

**Figure 2.**
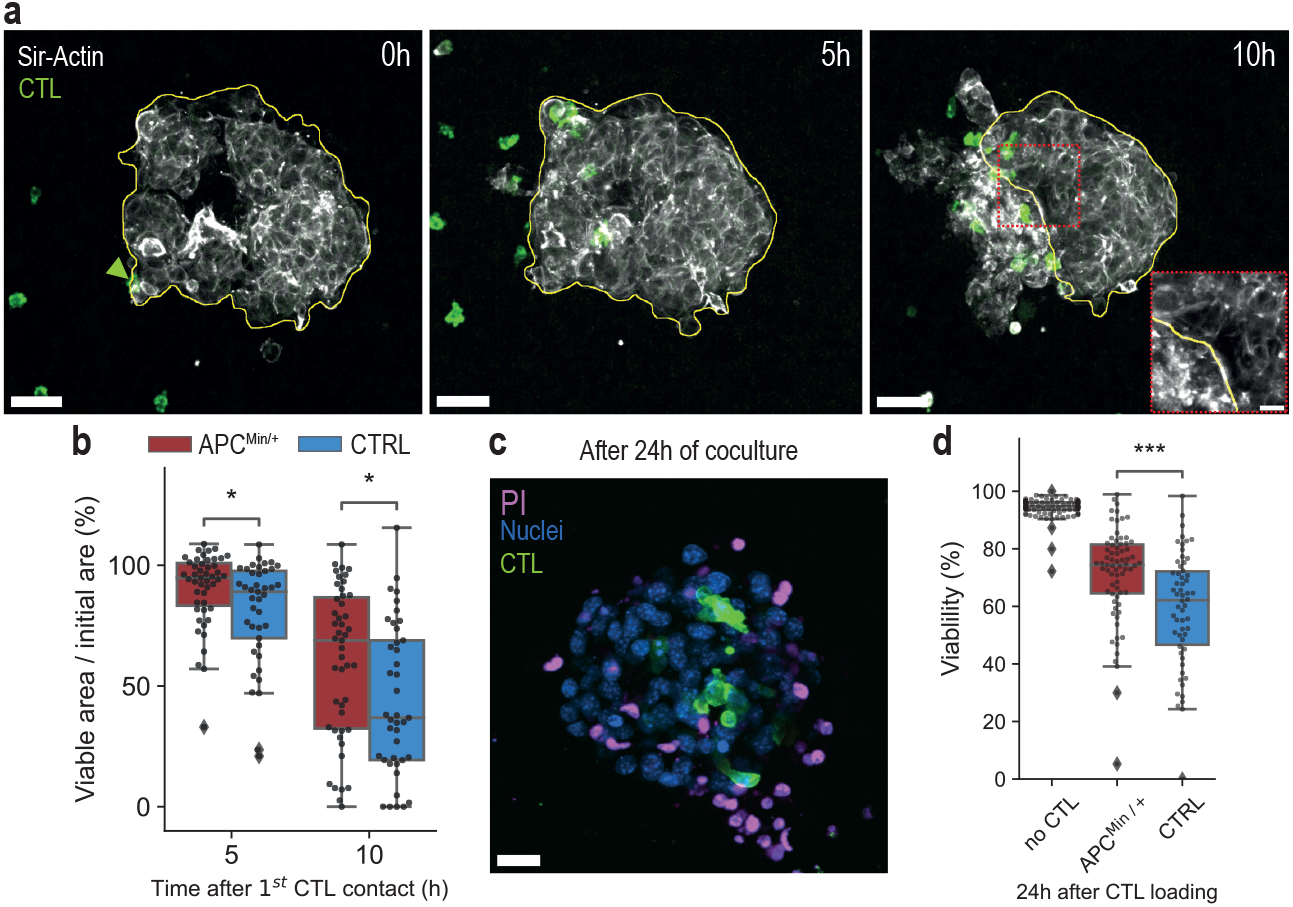
CTLs from APC^Min*/*+^ mice have a lower killing efficiency than control CTLs. **(a)** Representative images of spheroid destruction by CTLs. Time counted after the first CTL contact the spheroid (green arrow). The yellow contour is manually delimited. It corresponds to the part of the actin fiber network that remains organized. - Scale bars 50 µm. Inset: Zoom on the edge between living and dead regions to highlight differences in the actin fiber structure. - Scale bar 15 µm. **(b)** Quantification of spheroid destruction by CTLs measured as the portion of the initial area remaining alive, 5 h or 10 h after the first CTL contact (APC^Min*/*+^ : n=47 spheroids, control: n=41 spheroids). **(c)** Representative maximum intensity projection of a spheroid 24 h after CTLs addition – Scale bar 25µm. **(d)** Quantification of viability (using PI) after 24h of coculture (no CTL: n = 95 spheroids, APC^Min*/*+^ : n = 69 spheroids, control: n = 58 spheroids). - All statistical differences were calculated by the two-sided Mann-Whitney-Wilcoxon test.

The results shown in Fig. 2 highlight that, while both types of CTLs are able to induce B16-OVA death, APC^Min*/*+^ CTLs are less efficient at the level of the population. While the above measurements show a reduced ability of APC mutated cells to destroy cancer spheroids, it remains unclear which steps of the searching, accumulation or effective killing is limiting in the overall process. This question is subsequently addressed by dissecting the CTL behavior in the different periods of the experiment.

### APC^Min*/*+^ and control CTLs have similar ability to explore the 3D gel and accumulate on the tumor

The first step in the interaction between T cells and tumor cells is for the CTLs to find the spheroid and gather on it, which takes place in a random manner through spatial exploration [31]. For this reason we compared the migration dynamics of the two CTLs types in Matrigel, i.e. away from the spheroids, to investigate the role of their accumulation dynamics in the reduced ability to kill cancer cells. Tracking individual cells was possible due to an increased image acquisition rate, taking one image every three minutes (Fig. 3a and Movie S3). The measured mean-square displacement (MSD) was consistent with an exponent *α* = 1.4 for both cell types (Fig. 3b), in agreement with extensive literature on CTL migration statistics [31, 42, 43]. This similarity in MSD power indicates that the two CTL populations adopt a similar strategy to explore the space around the tumor and to find their targets.

**Figure 3.**
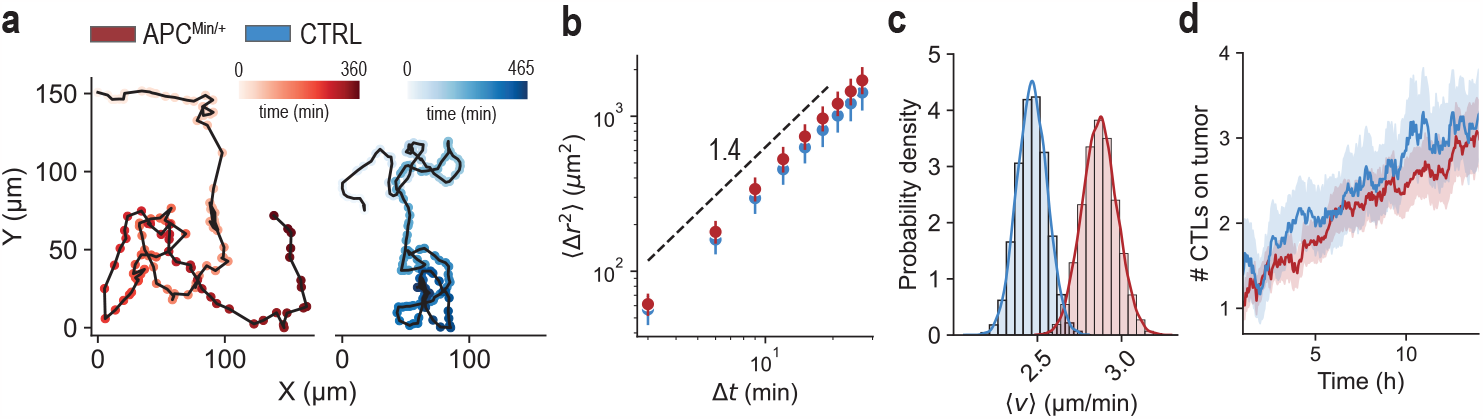
Mutated and control cells have a similar ability to explore the space in the Matrigel and to accumulate on tumor spheroids. **(a)** Representative trajectories of CTLs in the Matrigel matrix (not in contact with the spheroid). Color saturation indicates the time. Red: APC^Min*/*+^, blue: control. **(b)** Mean-square displacement (MSD) of CTL migration in Matrigel. Error bars represent 95 % confidence intervals. Dashed line corresponds to a slope of 1.4. **(c)** Probability densities of single cell average velocities (bootstrapped data - see Materials and Methods). Red: APC^Min*/*+^, blue: control. **(d)** Number of CTLs accumulating on the spheroids after the beginning of imaging. APC^Min*/*+^ (red): n = 51 spheroids. control (blue): n = 25 spheroids. - Error bars represent the s.e.m.

Comparing the average CTL velocity in the Matrigel, APC^Min*/*+^ CTLs migrate slightly faster than control cells (Fig. 3c). The difference, however, remains modest, with an increase in the average velocity of 16 %. APC^Min*/*+^ CTLs can therefore explore the gel and find the tumor as quickly as the control cells. Finally, measurement of the accumulation of CTLs on tumor spheroids over time showed that the two types of CTLs accumulate in the same way on the spheroids (Fig. 3d). This is consistent with the similarity in number of CTLs observed 5 h and 10 h after first contact in the previous experiment (SI Fig. S2c). Altogether, these measurements reveal no clear effect of APC mutation on the exploration of space by the CTLs around the spheroids and their accumulation on the target tumor.

Tracking of some CTLs during the transition from the Matrigel to the spheroid (SI Fig. S3a) revealed a sharp decrease in their velocity when coming into contact with cancer cells (SI Fig. S3b-c). This might be attributed to a change of migration mode of CTLs from migrating along Matrigel fibers in 3D around the tumor [44–47] to exploring the 2D surface of the spheroid. Notably, T cells are also able to penetrate inside the tumor spheroid as some cells can be found inside the aggregate by confocal microscopy (SI Fig. S3d-e - green arrows - and Movie S4).

### APC mutated CTLs migrate faster and engage less with the cancer cells than control CTLs

CTL dynamics on the spheroids can be quantified by tracking individual T cells migrating on the tumor (SI Fig. S4a). First, the MSDs of the trajectories show that APC^Min*/*+^ CTLs explore the spheroids faster than the control cells, although they have similar exponent *α* of the power law at short time scale (Fig. 4a and SI Fig. S4b). This means that the two T cell populations migrate with similar dynamics but that APC^Min*/*+^ CTLs have a larger prefactor in the MSD. While the average velocity in the gel was similar for both cell types, the APC^Min*/*+^ CTLs move at a larger speed on the surface of the spheroid, compared with the control cells (Fig. 4b).

**Figure 4.**
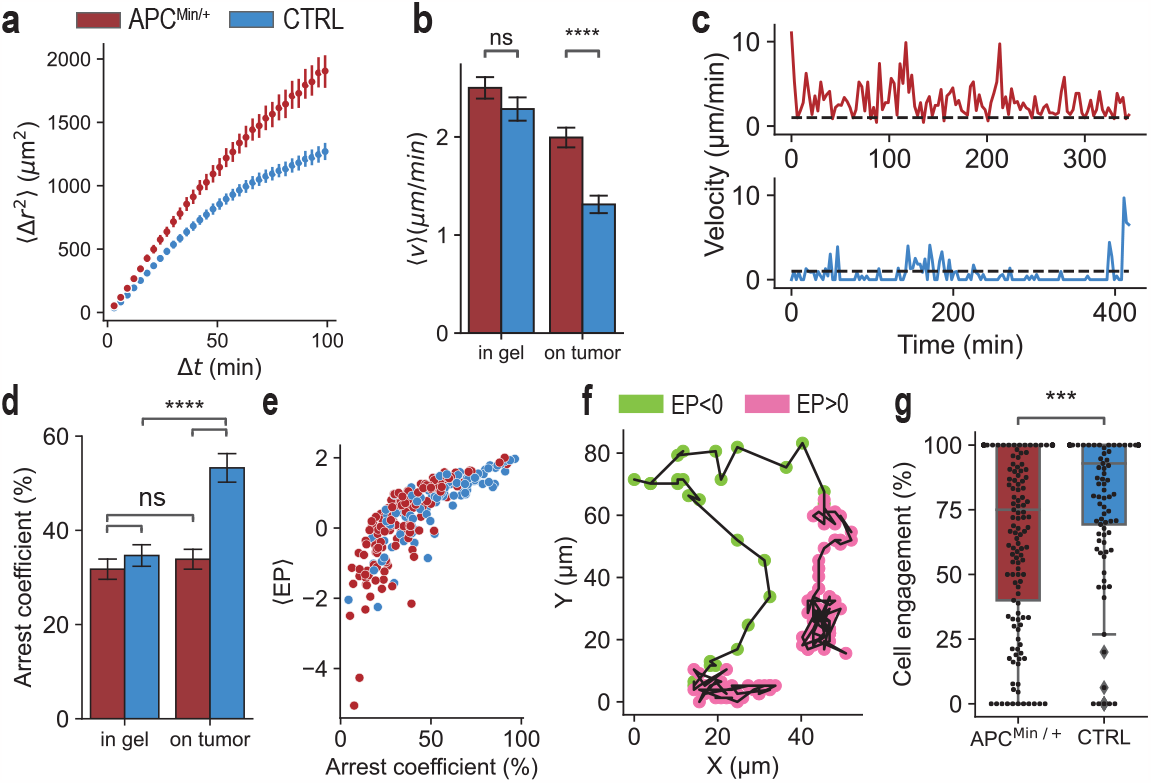
CTLs from APC^Min*/*+^ mice display modified migration on the spheroid and reduced arrest periods. **(a)** Mean-square displacement (MSD) of CTL migration on tumor spheroids. Error bars represent s.d. after bootstrapping data 100 times (APC^Min*/*+^ CTLs: n = 173, control CTLs: n = 106). **(b)** Single cell average velocity in the Matrigel and on the spheroid - average per droplet. **(c)** Representative velocity as a function of time for a control CTL (blue) and APC^Min*/*+^ CTL (red). **(d)** Single cell arrest coefficient in the Matrigel and on the spheroid. The arrest coefficient was defined as the percentage of time during which instantaneous velocity was <1 µm/min. **(b)-(d)** (in Matrigel: n = 45 droplets for control CTLs and n = 59 droplets for APC^Min*/*+^ CTLs. / on spheroid: n = 25 droplets for control CTLs and n = 32 droplets for APC^Min*/*+^ CTLs.) - Statistical differences were calculated by Welch’s t-test. - Error bars represent s.e.m. **(e)** Average engagement parameter along individual cell trajectories against arrest coefficient (APC^Min*/*+^ CTLs: n = 173, control CTLs: n = 106). **(f)** Representative trajectory of a cell on a tumor spheroid. Green regions correspond to *EP <* 0, when the cell is exploring the spheroid surface. Pink regions correspond to *EP >* 0, when the cell engage with the tumor locally. **(g)** Percentage of time spent in engaged phase (*EP >* 0) for individual cell trajectories for APC^Min*/*+^ and control CTLs (APC^Min*/*+^ CTLs: n = 173, control CTLs: n = 106). - Statistical differences were calculated by Welch’s t-test.

Further insight into the motility statistics was obtained by observing the time-dependent velocity of each cell. We find that the cells alternate between phases of high velocity and phases of arrest, as shown in Fig. 4c. By defining an arrested state as a period when the velocity is below 1 µm/min, the arrest statistics for each cell type could be compared. In particular, we measured the arrest coefficient, as the ratio of time spent in the arrested state vs. the motile state, for each track. The control cells displayed a large increase in the arrest coefficient after they transition from the gel to the spheroid, in contrast with the APC^Min*/*+^ cells that maintained the same value of the arrest coefficient in both environments. Furthermore, the duration of the arrest periods for the APC mutated cells was much shorter than for the control cells (see SI Fig. S4c).

Although the arrest coefficient is informative and often used to characterize immune cell dynamics [48, 49], it is nevertheless sensitive to instantaneous variations of velocities compounded by the uncertainty from the tracking process. Specifically, some cells can change their shape, displaying a high velocity but are localized and still induce killing (Movie S5). A more precise measure of the switching between exploration and engagement of CTLs was therefore computed from a combination of kinetic energy and radius of gyration of the cell trajectory (inspired from [50], see Materials and Methods, and Fig. S4d-i for details). We termed this novel parameter engagement parameter (*EP*) and used it to distinguish two different modes of migration. For negative values of *EP*, cells travel fast over large distances while for positive values of *EP*, cells densely explore a small region (SI Fig. S4g). The statistics of *EP* correlate well with the arrest coefficient, as expected from the physical interpretation of the two parameters (Fig. 4e). However, *EP* values allow the separation of individual trajectories into portions in which the cells are engaged with the tumor vs. portions in which they are traveling over large distances (Fig. 4f and SI Fig. S4h-i). By summarizing the relative periods of engagement vs. exploration of the CTLs on the spheroids, we observe a large difference in engagement between the mutant and control CTLs (Fig. 4g).

### APC^Min*/*+^ and control CTLs display high engagement with target cells prior to spheroid apoptosis

The difference in the ability of cells to arrest and engage with the tumor spheroid reveals a potential defect in CTL-tumor interaction due to the APC mutation. The main killing pathway of T cells relies on the formation of an immunological synapse with their target after finding the cognate antigen [39, 51–53] and this event leads to an arrest of the T cell [32, 47, 54, 55]. The lack of arrest and engagement observed above may therefore be related with reduced ability to form the immunological synapse and can explain the reduced cytotoxicity of the APC mutant. To link CTL activity to tumor cell death, caspase 3/7 activation was measured to identify cell apoptosis events [56], as CTLs are known to induce tumor cell apoptosis [31, 38]. In these experiments caspase 3/7 activation was recorded every 30 minutes, while CTL dynamics were imaged every 3 minutes, as above (Movie S6). The caspase 3/7 signal showed two distinct modes of activation. In some cases the activation of caspase 3/7 occurred in isolation for individual cells, while in other cases, we observed a cascade of several cells being activated in a localized region of the spheroid and in close succession (see Fig. 5a and Movie S6). Caspase 3/7 activation cascades were observed in 67 % of the spheroids and their existence correlated with high mortality of the aggregate after 24 h of coculture with CTLs, as measured by PI staining (SI Fig. S5a).

The existence of these localized killing events therefore provides a link between CTL dynamics and their ability to kill the cancer cells, by comparing the CTL engagement in these spatio-temporal windows with their overall dynamics. By looking backwards in time after a killing cascade occurred it was possible to estimate the nucleation time and location of each of these cascades, keeping in mind that some spheroids displayed several independent cascades at different times and locations. Since the caspase 3/7 signal results from a complex process that potentially involves several sub-lethal hits by the CTLs [16, 19, 20], we analyzed the CTL behavior in a 50 µm radius around the nucleation event and looking backwards in time, as sketched by the dotted circle in Fig. 5b. As CTLs can elongate over large distances (>40 µm, SI Fig. S5b) and caspase signal accumulates in a large region (SI Fig. S5a, Movie S6), 50 µm was chosen to account for T cells involvement in the spheroid destruction.

The number of individual APC^Min*/*+^ cells detected in the region, in a 4 h period before the caspase 3/7 signal nucleation, was larger than the control (Fig. 5c), which was not the result of a different number of CTLs detected on the spheroid (SI Fig. S5c). By measuring the average *EP* in the 4 h before caspase signal, APC^Min*/*+^ cells are less engaged than the control. However, a sharp transition to the engaged state can be observed approximately 2 h before killing initiation, with the average value of the *EP* of APC^Min*/*+^ cells significantly increasing and reaching a similar value as for the control cells (Fig. 5d). Only the realignment of CTL trajectories to the time of the caspase activation allows us to visualize this transition to engaged state (SI Fig. S5d). Therefore, the higher number of CTLs observed in the 4 hours before killing for APC^Min*/*+^ cells (Fig. 5c) can be explained by an overall lower engagement of APC^Min*/*+^ cells leading to a increase number of CTLs only passing through the region and not engaging properly. Moreover, the observed transition to high engagement observed before caspase 3/7 activation, allows us to link CTL dynamics with efficient tumor killing. As high T cell engagement precedes cancer cell death, the observed reduced engagement of APC^Min*/*+^ CTLs compared to the control (Fig. 4d,g) explains the observed reduction in cytotoxicity of APC^Min*/*+^ cells (Fig. 2b,d).

**Figure 5.**
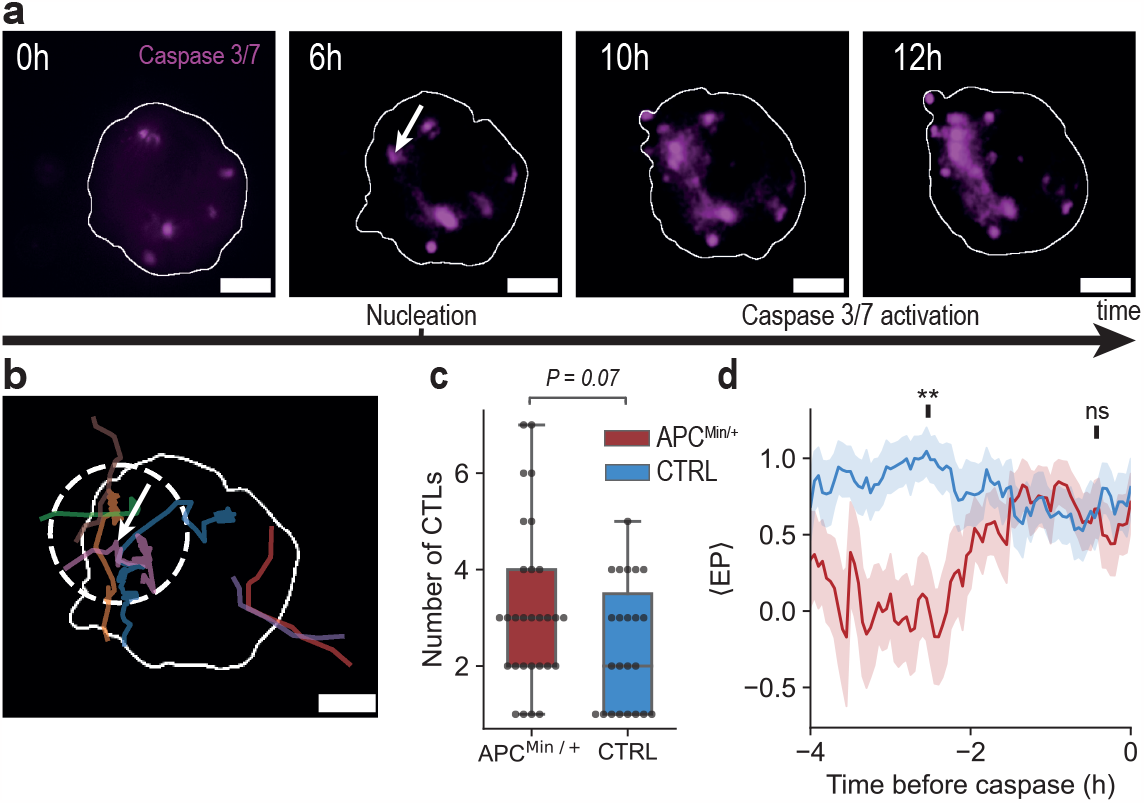
Localized apoptosis cascades associated with high engagement periods. **(a)** Representative images of caspase cascade and selection of nucleation initiation and position (white arrow). Time measured after the beginning of time-lapse imaging. The white line depicts the defined spheroid area. **(b)** Trajectories of CTLs before the observation of caspase signal. Each colored line corresponds to one CTL. The white arrow points to the region of caspase cascade nucleation. The dotted-line circle corresponds to a radius of 50 µm. All scale bars 50 µm. **(c)** Number of CTLs staying more than 15 min in the dotted circle in Fig. 5b in the 4 h window before the nucleation event. Statistical difference was calculated by the two-sided Mann-Whitney-Wilcoxon test (n = 29 events for APC^Min*/*+^ CTLs and n = 23 events for control CTLs). **(d)** Average engagement parameter of CTLs before initiation of caspase 3/7 activation (average per event *±* s.e.m). Statistical difference was calculated by Welch’s t-test 2.5 h and 0.5 h before caspase 3/7 signal. (n = 29 events for APC^Min*/*+^ CTLs and n = 23 events for control CTLs).

## DISCUSSION

The results shown here provide the first clear evidence that APC mutated CTLs have a reduced ability to destroy 3D tumor spheroids. Detailed analysis of the CTL trajectories shows that this reduction of killing efficiency is not due to an impairment of the motility in the gel nor accumulation of the CTLs, which were similar between the APC^Min*/*+^ and the control cells. Instead, the APC mutation is found to strongly reduce the engagement of the CTLs with the spheroid after contact has been made. Although we do not focus on the molecular aspects of these cell-cell interactions, these results are consistent with data from single-cell experiments on APC-silenced or FAP patient-derived T cells: A reduced mechanical stability of the immunological synapse [12] and an increase of the migration ability in collagen-coated channels [13]. The reduction in the CTL engagement and the reduced ability of the APC^Min*/*+^ cells to destroy the spheroids can both be due to the impaired capability to form a stable immunological synapse. Compared with the single-cell experiments however, the current results integrate, in a three dimensional setting, the complete chain of events that lead from the introduction of the T cells in the vicinity of the spheroids to its destruction. Importantly, a novelty of our approach is that the trajectory analysis provides detailed insight about the net impact of the reduced strength of the immunological synapse in realistic cell-cell interactions.

The experiments shown here can distinguish differences in the CTL-cancer interactions for even closely related cell types. The ability to identify subtle differences is enabled first by tracking a large number of trajectories in many independent droplets, each of which corresponds to an independent replicate of the experiment. Second, the dynamic multiscale cytometry [31, 35], which is constructed by linking the global outcome with measurements of many individual trajectories, allows us to divide the dynamics into separate periods that are treated independently. Here, a new metric (*EP*) is introduced to describe the individual trajectories. It has a reduced value in the APC^Min*/*+^ cells on average, in line with a reduced cytotoxicity. More interestingly, by focusing on key events in target cells (caspase 3/7 initial activation in a small region) and aligning the T cell behaviors to these precise spatio-temporal windows, the value of *EP* for APC^Min*/*+^ CTLs transitions from low values to reach the same value as the control cells in those windows.

From an applied bio-engineering point of view, the approach developed here can provide a robust platform to test the quality of CTLs in the context of adoptive cell therapy. Since all T cells in the droplets must go through a similar process of searching, accumulation, antigen recognition, and killing, each of these stages can be evaluated independently to identify possible deficiencies in the probed cells. The droplet-based approach offers specific advantages as it is based on a standardized experimental protocol in which a well controlled number of immune cells is added at a defined moment. Since the distances traveled and the size of spheroids are all reproducible between experiments, it is straight forward to compare the efficiency of each individual step in each of the experiments. As a result, analysis of these experiments can yield far greater details of CTL cytotoxicity than the current state of the art.

## MATERIALS AND METHODS

### Tumor cells

B16-Ovalbumin peptide (residues 257–264) expressing cell-lines (B16-OVA) were maintained in RPMI 1640 (Fischer Scientific 12027599) media containing 10% FBS and 1% penicillin-streptomycin antibiotics (Fischer Scientific – 10378016) at 5 % CO2 and 37°C. B16.F0OVA (B16-OVA) melanoma cells were kindly provided by Claude Leclerc. All experiments were conducted at a passage number smaller than 35.

### Mice, T lymphocyte isolation and cell culture

Mice were housed and bred under pathogen-free conditions in the Central Animal Facility of Institut Pasteur. The protocols used have been approved by the Ethical Committee for Animal Experimentation of the Institut Pasteur and by the French Ministry of Research (authorization nr. 15407-2018060518151060). Heterozygous C57BL/6J-ApcMin mice (APC^Min*/*+^) [8] were crossed to UBC-GFP Rag1^*−/−*^OT-I TCR transgenic mice (kindly provided by Dr. P. Bousso, Institut Pasteur [57]). APC^+*/*+^ and APC^Min*/*+^ littermates were sacrificed at 8–12 weeks of age and spleens were removed and homogenized through a 70 µm filter. To stimulate T cells and generate CTLs, cell suspensions were washed twice in culture medium (RPMI-1640 + GlutaMAX-I (Gibco) supplemented with 10 % FBS, 1 mM sodium pyruvate and nonessential amino acids, 50 µM 2-ME, 10 mM HEPES, and 1 % penicillin–streptomycin (v/v)), and then split in two aliquots: one third was pulsed with 50 µM OVA (257-264) peptide (Anaspec AS-60193-1) for 2 h at +37°C, then it was mixed to the remaining splenocytes and incubated for 48-72 h at +37°C in a 5 % CO2 incubator. Cells were then counted, adjusted at 5*×*10^5^ cells/ml and cultured 2-5 additional days in cell culture medium supplemented with 20 ng/ml IL-2 (Miltenyi Biotec) before being used in the experiments.

### Microfabrication

The microfluidic mold to produce PDMS-based microfluidic device was 3D printed using HTM 140 V2 resin with Digital Light Processing printer (DLP MicroPlus EnvisionTec). Chips were produced and treated as described in Refs. [33, 35, 58]. In short, poly(dimethylsiloxane) (PDMS, SYLGARD 184, Dow Corning, 1:10 (w/w) ratio of curing agent to bulk material) was poured over the mold and cured for 2 h at 80°C. The top of the chip was sealed on a glass slide after plasma treatment. The chips were filled two times with Novec Surface Modifier (3M, Paris, France), a fluoropolymer coating agent, for 20 min at 110°C on a hot plate.

### Microfluidic setup

The loading of microfluidic chips was done according to the protocol described in Ref. [31]. More precisely, chips and oil were cooled down at -20°C to prevent Matrigel polymerization. The first droplet contained a concentration of 0.6 *×* 10^6^ cells/mL B16-OVA melanoma cells encapsulated in 0.7 µL of RPMI media and Corning Matrigel (Dutscher Dominique – 354234) at a concentration of 1.8 mg/mL (20 % of Matrigel at 8.9 mg/mL). The oil phase around the droplets was composed of fluorinated-FC40 (3M) oil mixed with 2 %(v/v) FluoroSurfactant (Ran Biotechnologies). Chips were placed at 4°C for 20 min after loading to allow cell sedimentation without Matrigel polymerization and avoid the formation of multiple spheroids. Chips were then incubated at 37°C for 24 h. Second droplets, containing RPMI media and OT-I-CD8^+^GFP cells at concentrations varying from 1 to 2 *×* 10^6^ cells/mL in a volume around

0.25 µL, were trapped in the draft-shape traps (Fig. 1a,e, SI Fig. S1a). Droplet fusion was ensured by the addition of 20 % (v/v) of 1H,1H,2H,2H-perfluoro-1-octanol (PFO) (Sigma-Aldrich) dissolved in NovecTM-7500 Engineered Fluid (3M). After fusion, PFO was removed from the chamber by addition of 1.5 mL of pure FC40 oil in the chamber. Chips were flipped upside-down during 30 min to facilitate T cells entry in the Matrigel phase before being imaged.

### Microscopy

Confocal images were acquired using spinning disk confocal microscope (Nikon Ti2 + Yokogawa) with a 20x 0.7 NA air objective lens (Nikon Inc.). Epifluorescence and bright field images were captured using a Nikon Ti2 motorized epifluorescence microscope with a 20x objective lens. The illumination was produced by a Lumencor LED light source, and the images were captured by a Hamamatsu C13440-20CU SCMOS camera. Raw data collection was done using imaging software Nikon Elements (version 5.11.01, Build 1367).

### Proliferation measurement

B16-OVA proliferation was assessed with a BrdU staining kit (eBioscience). B16-OVA spheroids were formed as described above. 24 h after cell loading, 200 µL of a solution containing 50 % (v/v) of 1H,1H,2H,2H-perfluoro-1-octanol (PFO) (Sigma-Aldrich) dissolved in NovecTM-7500 Engineered Fluid (3M) was injected in the chip to destabilize the droplet interface and allow the BrdU solution to diffuse inside the droplet of polymerized Matrigel. Immediately after, the PFO was replaced by 0.5 mL of a PBS (Sigma-Aldrich - D8537) solution containing 50 µM of BrdU (eBioscience - 00-4425-10). The chip was incubated at 37°C during 5 h. The BrdU solution was then washed with 0.5 mL of PBS. 0.5 mL of fixation solution, made up of 1/4 of BrdU staining buffer concentrate (eBioscience - 00-5515-43) and 3/4 of Fixation/Permeabilization Diluent (eBioscience - 00-5223-56), was then introduced in the chip. The chip was incubated overnight at 4°C on a rocker for mixing. The fixation solution was removed using 0.5 mL of PBS. Flow cytometry buffer was prepared diluting 1 % of FBS in PBS. DNase I (eBioscience - 00-4440-51A) was mixed in Flow cytometry buffer at 30 % and injected in the chip (0.5 mL). The chip was incubated at 37°C for 1 h. DNase I solution was washed using 0.5 mL of Flow cytometry buffer. 0.5 mL of Flow cytometry buffer mixed with an Alexa-488 conjugated anti-BrdU antibody (eBioscience - 11-5071-42) at 1/100 and NucBlue fixed cells (Fischer scientific - R37606) (1 droplet in 0.5 mL) was injected into the chip and incubated at room temperature for 4 h in the dark. 0.5 mL of PBS was used to wash the solution. Samples were imaged using confocal microscopy. Z-stack images every 10 µm were acquired over a range of 100 µm. Quantification was performed with ImageJ software. Z-intensity maximum projection was applied and the percentage of positive cells was assessed by computing the ratio of the BrdU positive area over the nuclei positive area.

### Viability and apoptosis measurement

For viability measurements, B16-OVA spheroids were formed as described above. 24 h and 48 h after cell loading, 200 µL of a solution containing 50 % (v/v) of 1H,1H,2H,2H-perfluoro-1-octanol (PFO) (Sigma-Aldrich) dissolved in NovecTM-7500 Engineered Fluid (3M) was injected in the chip. Immediately after, 300 µL of a solution containing RPMI media, Propidium Iodide (PI) (Sigma - P4864) at a concentration of 3 µM and NucBlue live cells (Fischer scientific - R37605) (2 droplets in 1 mL) was introduced in the chip and incubated 4 h at 37°C. Samples were imaged using confocal microscope. Z-stack images every 10 µm were acquired over a range of 100 µm. Quantification was performed with ImageJ software. Z-intensity maximum projection was applied and the percentage of positive cells was assessed by computing the ratio of the PI positive area over the nuclei positive area. For apoptosis measurements, B16-OVA spheroids were formed as described above and caspase 3/7 Red Apoptosis Assay Reagent (Essen Bioscience - 4704) was added in the initial media at a concentration of 2 µM. Samples were imaged using epifluorescence microscope. Quantification was performed with ImageJ software. The percentage of positive cells was assessed by computing the ratio of the caspase positive area over the spheroid area.

### Spheroid reduction

Spheroids were formed and CTLs were introduced as described in Microfluidic setup. Actin fibers were stained adding Sir-Actin and Verapamil (Spirochrome Cytoskeleton Kit - SC006) in the primary droplet media at respectively 1 µM and 10 µM. Time lapse acquisition was performed with confocal and bright field microscopy. Z-stack images every 10 µm over 100 µm were acquired every 30 min over 15 to 20 h. The time of CTL contact and the number of CTLs on the spheroid were counted manually. The portion of intact actin was manually outlined using ImageJ.

### CTL dynamics

Spheroids were formed and CTLs were introduced as described in Microfluidic setup. To be able to track the cells over time, images were acquired every 3 min over 15 to 20 h. CTLs were tracked in the Matrigel using Track-Mate software [59] in ImageJ. Because of overlapping trajectories, CTLs were manually tracked on the spheroid using ImageJ. Only trajectories longer than 30 min were considered. To remove dead cells from the Matrigel, cells moving less than 15 µm in a 30 min interval were not considered. Spheroids were segmented using ImageJ. CTL positions were overlapped with spheroid masks to compute distance between tumor and CTLs. Only cells at a distance > 50 µm from the tumor spheroid were considered for trajectories in the Matrigel.

### Spheroid apoptosis

Caspase 3/7 Red Apoptosis Assay Reagent (Essen Bioscience - 4704) was added to the primary droplet medium at a concentration of 2 µM. Images were acquired every 30 min.

### Bootstrapping

In order to concatenate data from several large datasets with different number of points, we used bootstrapping to minimize bias. This methodology was applied for Fig. 3c and Fig. 4a. In Fig. 3c, 100 trajectories from each of three independent experiments were randomly selected and pooled together. The average velocity was computed over 200 trajectories randomly selected from these 300 trajectories. The procedure was repeated 10.000 times and the probability density of average velocities is plotted. In Fig. 4a, 15 trajectories from each of three independent experiments were randomly selected and pooled together. The average MSD was computed over 20 trajectories randomly selected from these 45 trajectories. The procedure was repeated 100 times. Average values and standard deviations are shown over these 100 repetitions.

### Engagement parameter computation

To distinguish between periods of local motion and periods of fast exploration, we cut trajectories into segments of one hour length. This avoids bias due to differences in length of trajectories. The engagement parameter (*EP*) was computed in several steps. First, the gyration radius g and kinetic energy E were computed as follows: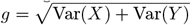 where Var(*X*) (resp. Var(*Y*)) is the variance of the *x* component (resp. *y* component) of the cell position and *E* is the average velocity square. These two metrics correlate (SI Fig. S4d) and hence, as a second step, the first component of the principal component analysis (*P*) was used as it describes 88 % of the variance in the data. In a third step, we define *EP* to distinguish between the engaged state, when the cell explore locally the spheroid (*EP >* 0), and the exploring state when the cell move fast (*EP <* 0). To this end, *EP* was computed from *P* by *EP* =*− P* + *C* where *C* is a constant, defined in such a way that the probabilities of the two states determined by fitting a Gaussian Mixture Model intersect at 0 (SI Fig. S4e-f). Example of pieces belonging to each state (randomly selected from the dataset) are shown in SI Fig. S4g. To distinguish engaged and exploration phases of single trajectories, *EP* was computed on a one hour rolling window over the trajectories. Engaged phases (*EP >* 0) and exploration phases (*EP <* 0) can be visualized on single cell trajectories (Fig. 4f and SI Fig. S4h-i).

## Data analysis

All data analysis and statistical tests were performed using Python (ns.0.05<P<1, ^*^P<0.05,^**^P<0.01,^***^P<0.001 and ^****^P<0.0001).

## AUTHORS’ CONTRIBUTIONS

V.B., V.D.B. A.A. and C.N.B. designed experiments. V.B., V.D.B. and M.Mad. performed experiments. C.C. and M.Mas. performed preliminary experiments. V.B. performed image analysis. V.B. and E.M. analyzed data. N.V.Q. and V.B. designed microfluidics. A.A. and C.N.B. supervised the research. V.B. wrote the original draft. V.B. and C.N.B. wrote, reviewed and edited the manuscript. All authors discussed the results and commented on the manuscript.

## ACKNOWLEDGMENTS

The authors acknowledge insightful discussions with Sébastien Sart, Andrey Aristov and all members of the Baroud and Alcover teams. The authors acknowledge the support of the Biomaterials and Microfluidics platform as well as FabLab at Institut Pasteur. Erik Maikranz has received funding from the European Union’s Horizon 2020 research and innovation programme under the Marie Skłodowska-Curie grant agreement N°899987. M. Madec was funded by an Institut Pasteur Cancer Initiative Masters scholarship, M. Mastrogiovanni has received a scholar of the Pasteur Paris University International Doctoral Program and was supported by the Institut Pasteur, the European Union Horizon 2020 Research and Innovation Programme under the Marie Sklodowska-Curie grant agreement 665807 (COFUND-PASTEURDOC) and La Ligue Contre Le Cancer-Allocation doctorale 4ème annee de thèse. This work has benefited from state financial aid, managed by the Agence Nationale de Recherche under the investment program integrated into France 2030, project reference ANR-21-RHUS-0003.

## SUPPLEMENTARY MOVIES

**Movie S1** - Illustration of droplet fusion and oil exchange in a microfluidic chip. For droplet fusion, the first droplet contains a spheroid of B16-OVA cells in medium mixed with 20 % of Matrigel and the second droplet contain CTLs. For oil exchange, the oil phase is replaced by medium phase that is merged with the main droplet. - Time in seconds, scale bar 200µm.

**Movie S2** - Time-lapse of B16-OVA spheroid under CTL attack using confocal microscopy. Images were acquired every 30 minutes. Actin fibers are stained in grey and CTLs are GFP. Two representative spheroids are shown. - Time in hours, scale bar 50µm.

**Movie S3** - Time-lapse of CTL migration in the Matrigel and on a spheroid. Images were acquired every 3 minutes. Bright field image and GFP channel are shown as well as CTL tracks and spheroid boundary. - Time in minutes, scale bar 50µm.

**Movie S4** - Time-lapse showing a CTL (GFP) infiltrating a spheroid using confocal imaging. The movie only shows the top of the spheroid with actin fibers in grey (sir-actin) until 4h30. Green arrows show an individual CTL. z-stacks every 10µm are shown every 30min. - Time in hours, scale bar 50µm.

**Movie S5** - Time-lapse showing a CTL (GFP) moving fast but very locally. Bright field imaging shows the spheroid and GFP signal is the T cell. - Time in minutes, scale bar 50µm.

**Movie S6** - Time-lapse showing a representative example of an apoptosis cascade on a tumor spheroids. Green signal corresponds to T cells and magenta to Caspase3/7 activation. White arrows highlight the beginning of the cascade of killing events. Images were acquired every 3 minutes for the CTLs and every 30 minutes for the Caspase3/7. - Time in minutes, scale bar 50µm.

---

**Figure S1.**
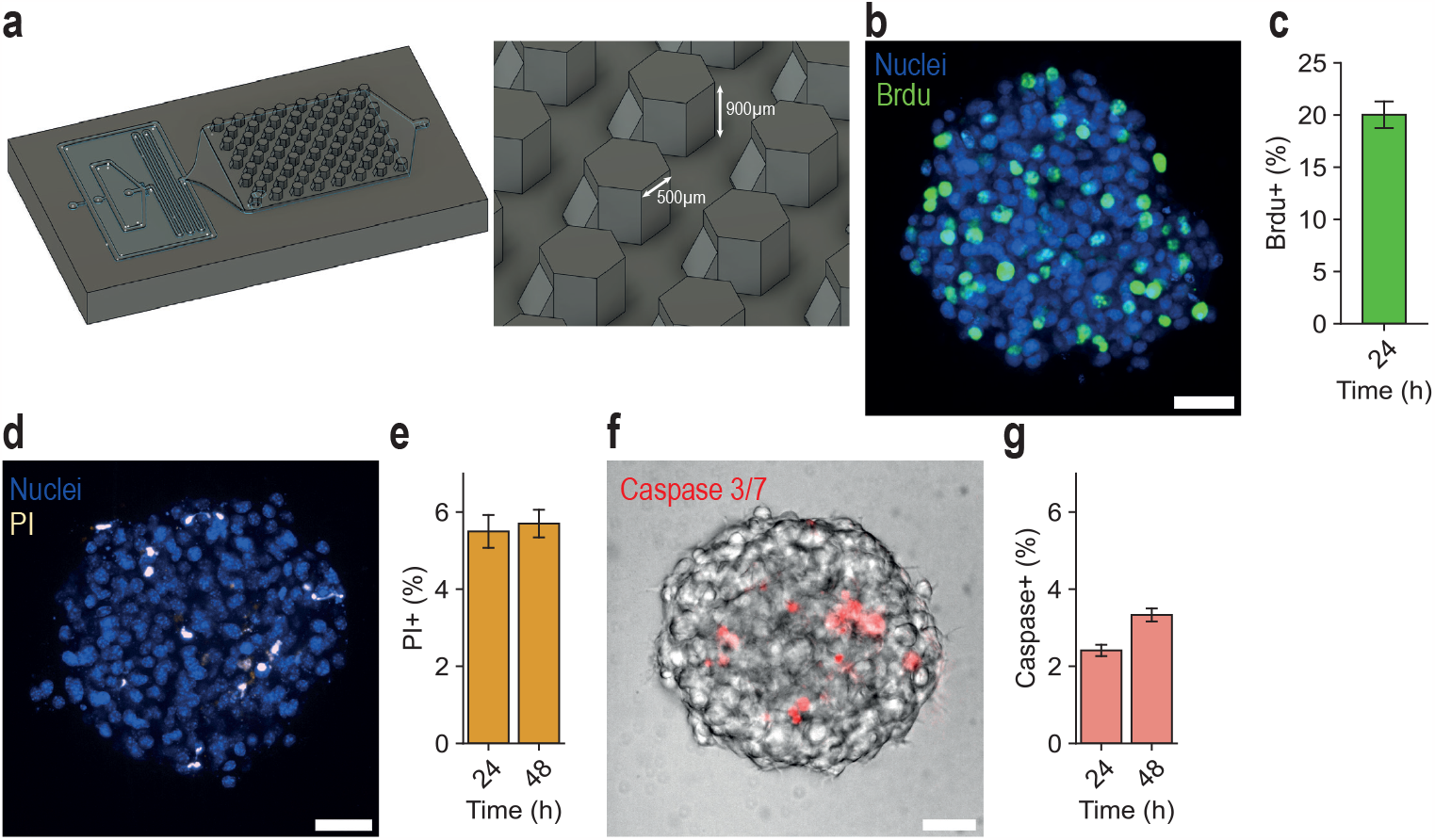
Microfluidic chip design and characterization of B16-OVA spheroids behavior in microfluidic chip. **(a)** 3D sketch and dimensions of 3D-printed microfluidic chip mold design. **(b)-(c)** BrdU staining of proliferating cells after 24 h in the droplet (n = 69 spheroids). **(d-e)** Propidium iodide (PI) staining shows high viability after 48 h (n = 45 and 95 spheroids). **(f-g)** Caspase 3/7 staining reveals low number of apoptotic cells after 48 h (n = 100 spheroids). - All scale bars 50 µm, error bars represent s.e.m.

**Figure S2.**
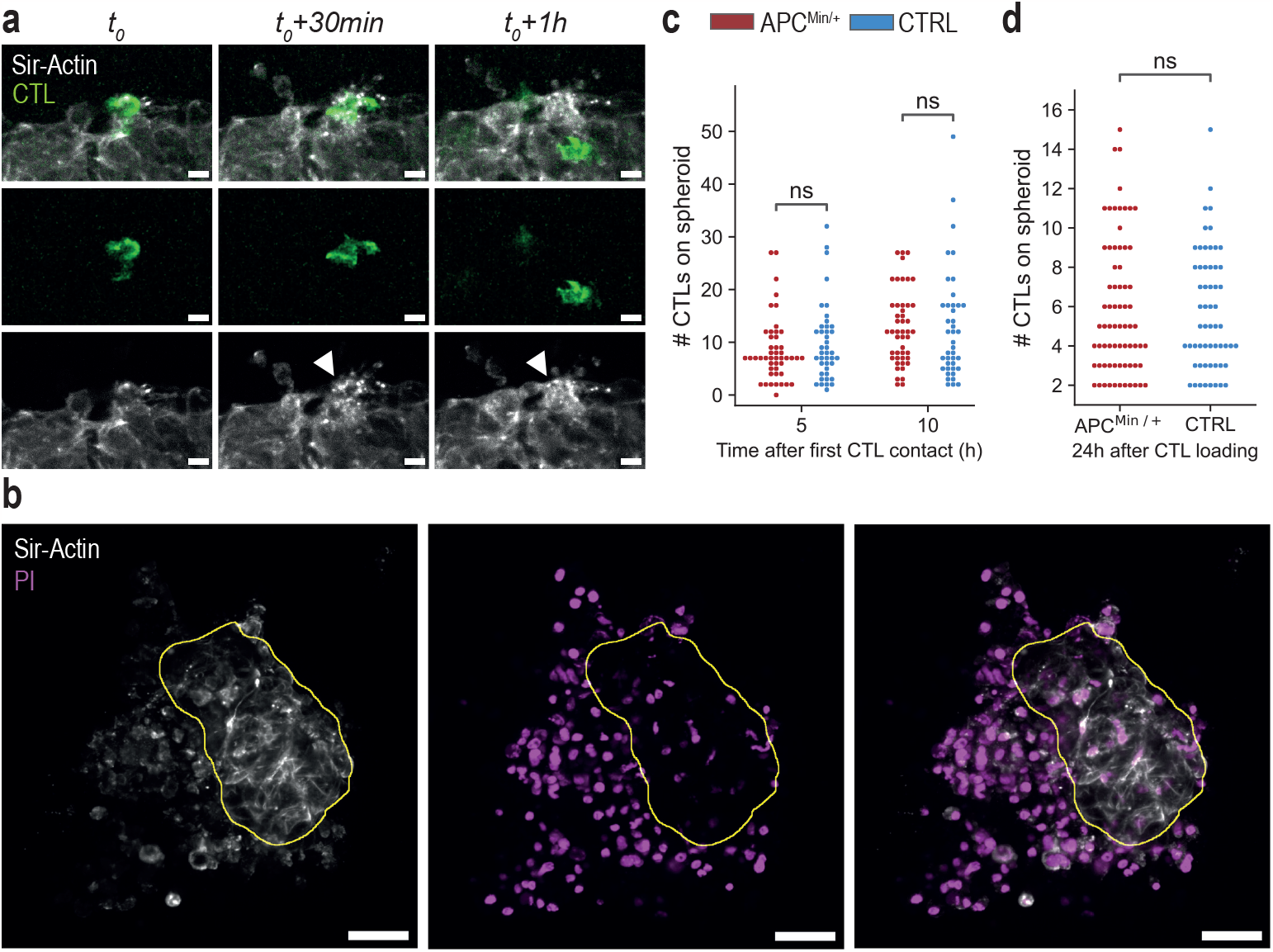
Actin cytoskeleton disorganization and absence of bias in CTL accumulation. **(a)** Representative images of T cell hit on actin fibers - top: Both channels, middle: CTL, bottom: Sir-Actin. White arrows show actin fragmentation. Scale bars 15 µm. **(b)** Representative images of a spheroid 24h after CTLs addition in the droplet stained with Propidium Iodide (PI) for death – left: Sir-Actin, middle: PI, right: Both channels. The region of organized actin fibers correlates with low spheroid death. The area circled in yellow is considered as a proxy for the part of the spheroid remaining alive. Fig. 1a and Fig. S2b show the same spheroid. Scale bars 50 µm. **(c)** Number of CTLs on the spheroid 5 h and 10 h after first contact (relative to Fig. 2b). **(d)** Number of CTLs on the spheroids after 24 h (relative to Fig. 2d). - Statistical differences were calculated using the two-sided Mann-Whitney-Wilcoxon test.

**Figure S3.**
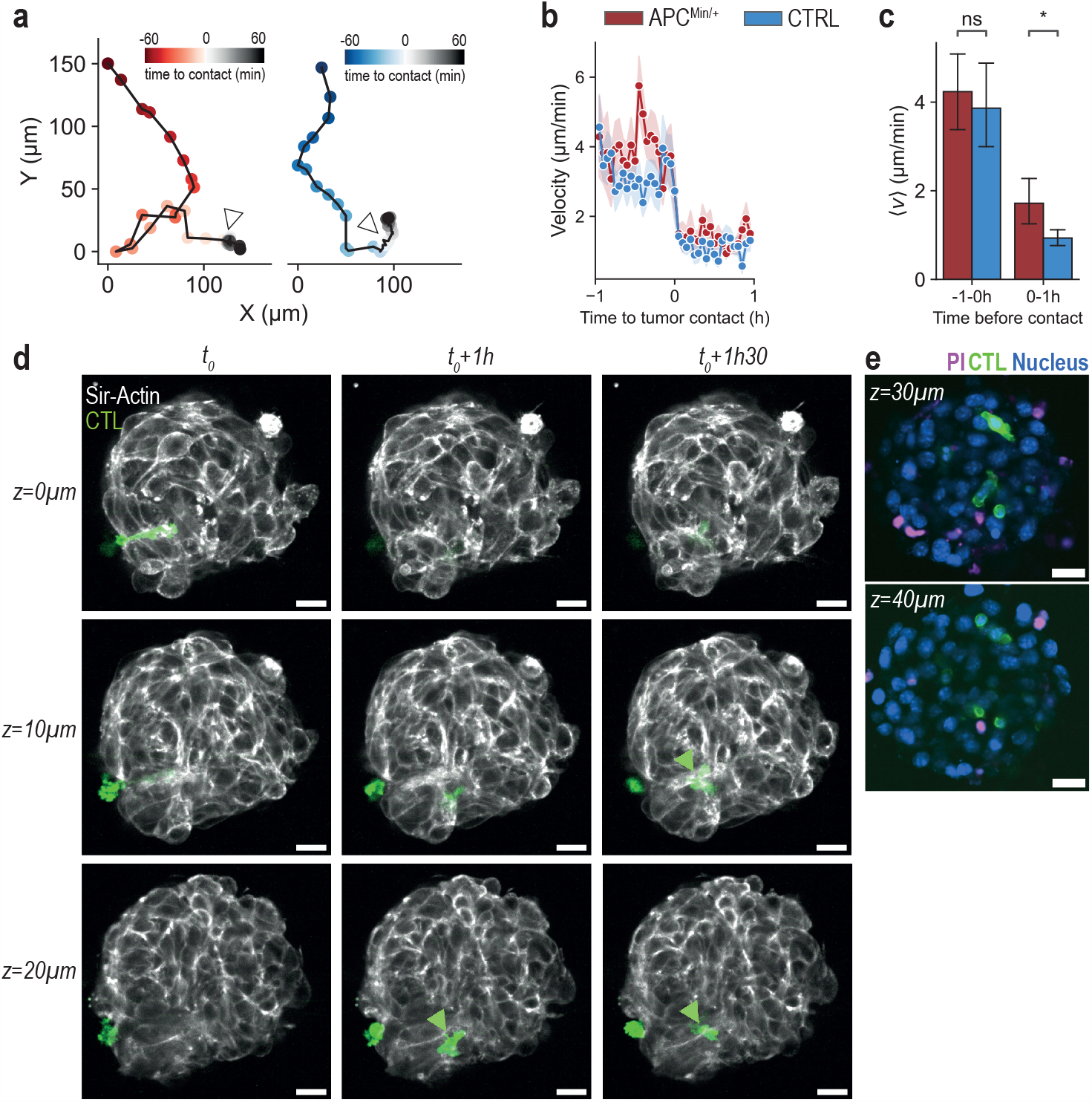
Transition from Matrigel to tumor and CTL penetration inside the spheroid. **(a)** Example of CTLs trajectories when finding the tumor. Color gradients correspond to time. Blue (in the Matrigel) to gray (on the spheroid): control CTL, Red (in the Matrigel) to gray (on the spheroid): APC^Min*/*+^ CTL. **(b-c)** Instantaneous and average T cell velocity 1 h before and after contact with the tumor spheroid (n = 8 individual cells control and n = 8 APC^Min*/*+^). Statistical differences were calculated using Welch’s t-test. Error bars represent s.e.m. **(d)** Dynamics of a T cell going inside a tumor aggregate (representative images). Green arrow highlight a T cell penetrating. Z increases when going deeper into the spheroid. **(e)** Representative images of a spheroid 24 h after CTLs addition. Some CTLs can penetrate – All scale bars 25 µm.

**Figure S4.**
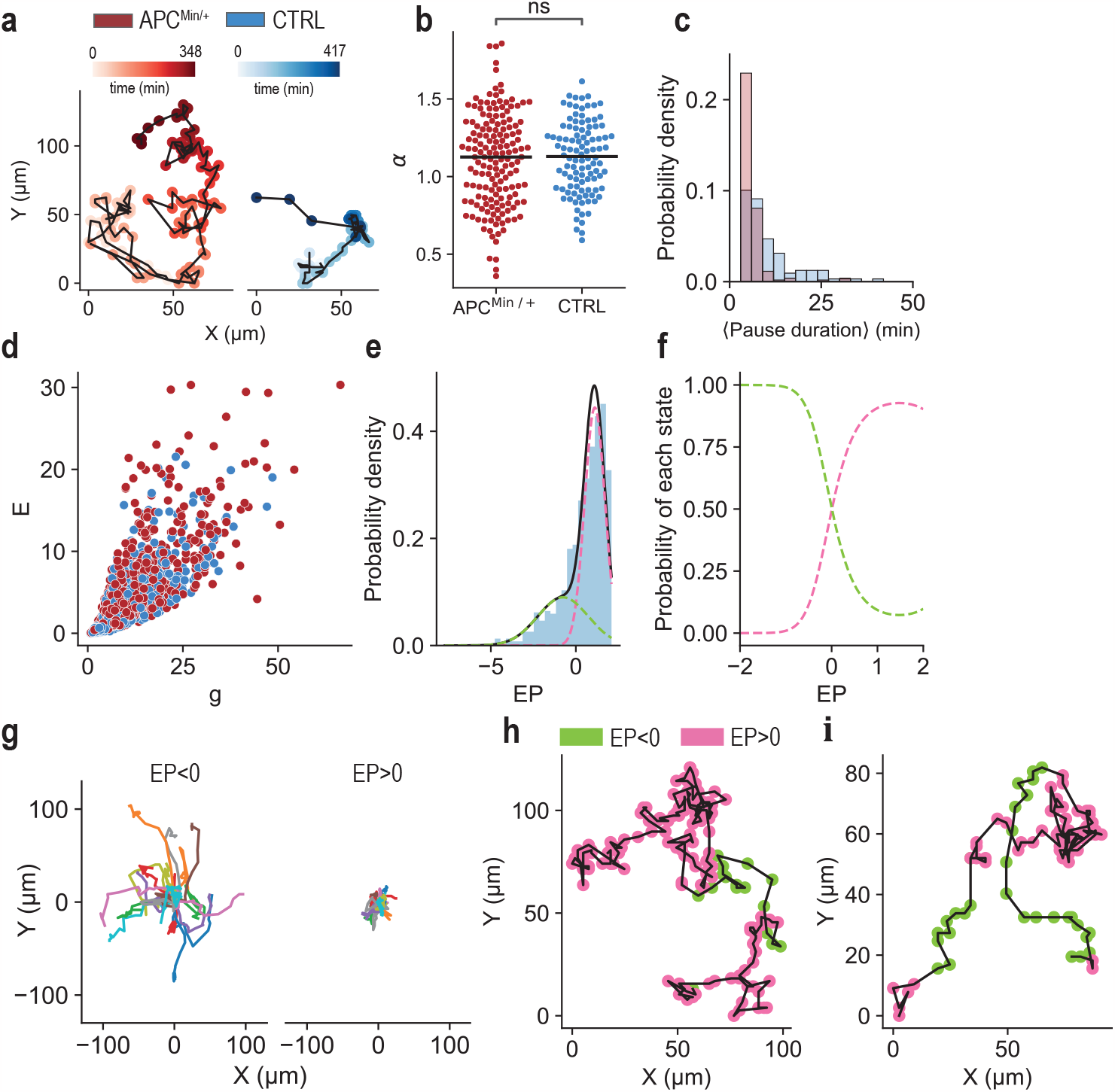
Complements to Fig. 4. **(a)** Representative trajectories of CTLs on the spheroids. Color gradients correspond to time. Blue: control CTL, Red: APC^Min*/*+^ CTL. (Same cells in Fig. 4c). **(b)** Coefficient of persistence *α*, determined by fitting a straight line to the MSD of individual cells in a log-log plot (only for ∆*t* < 24 min and if R^2^ > 0.8 for the line fit). Statistical differences were calculated by the two-sided Mann-Whitney-Wilcoxon test. Each point corresponds to one CTL (n = 106 for control CTLs and n = 173 for APC^Min*/*+^ CTLs). **(c)** Distributions of average pausing times on spheroids for individual CTLs. Pausing time is defined as the consecutive time with an instantaneous velocity <1 µm/min. **(d)** Kinetic energy against gyration radius for 1 h pieces of trajectory (n = 544 for control CTLs and n = 643 for APC^Min*/*+^ CTLs). **(e)** Engagement parameter density for 1 h pieces of trajectory (n = 1187). Dashed lines correspond to a fitted 2-states Gaussian Mixture Model. **(f)** Probability of each states of the fitted 2-states Gaussian Mixture Model. **(g)** 20 pieces of trajectory of 1 h (10 APC^Min*/*+^ and 10 control CTLs) in the high engaged (*EP >* 0) and low engaged (*EP <* 0) states (randomly selected). **(h)-(i)** Representative trajectories of a cell on a spheroid tumor. Green regions correspond to *EP <* 0, when the cell is moving fast and straight and pink regions correspond to *EP >* 0, when the cell explore locally the tumor.

**Figure S5.**
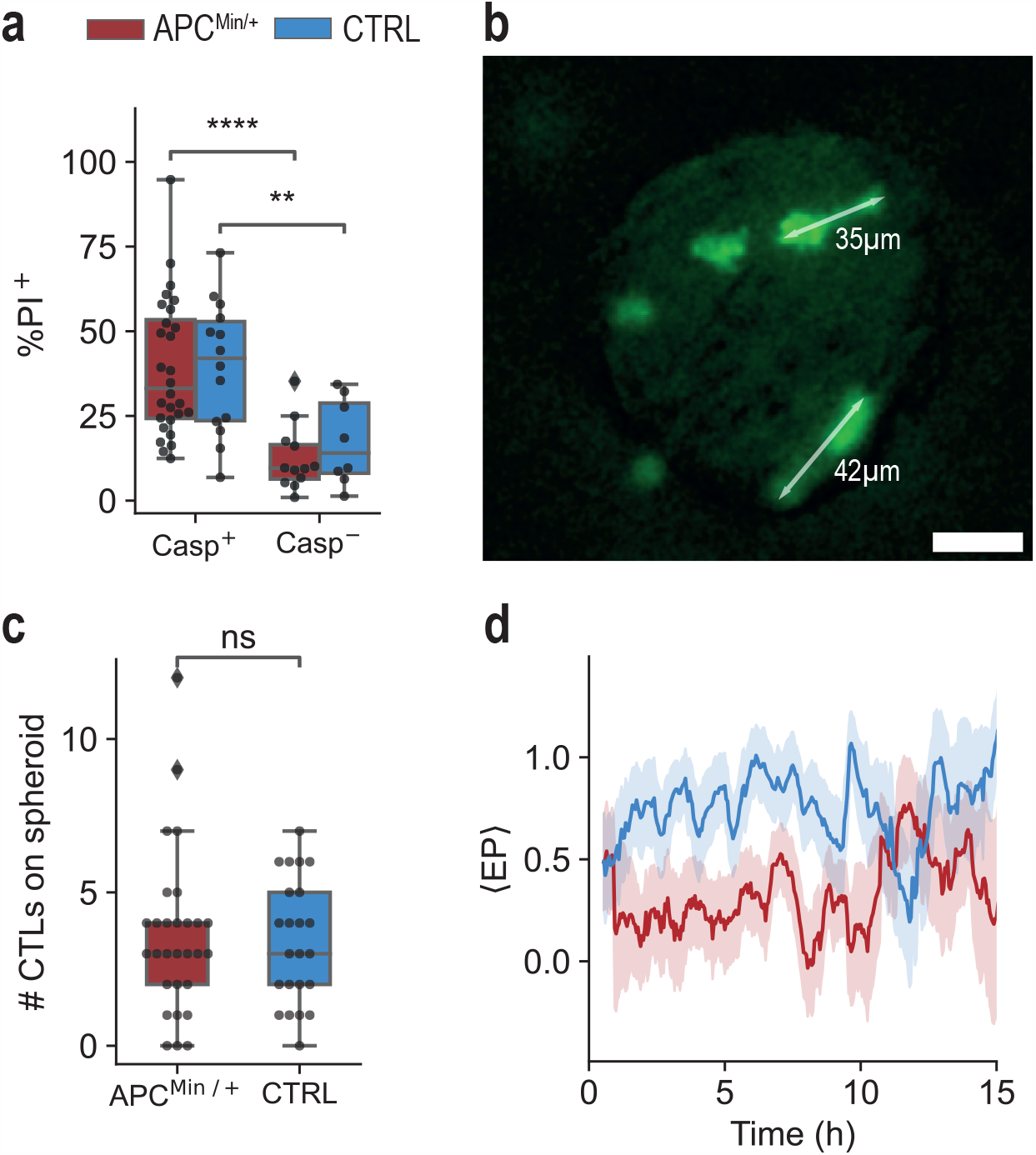
Complements to Fig. 5. **(a)** Percentage of dead cells (stained with PI) at 24 h of coculture according to the detection of caspase signal during time-lapse microscopy. Statistical differences were calculated using the two-sided Mann-Whitney-Wilcoxon test (n = 40 spheroids for APC^Min*/*+^ CTLs and n = 22 spheroids for control CTLs). **(b)** Representative image showing CTL elongation. - Scale bar 30 µm. **(c)** Number of CTLs on the spheroid at the time of caspase nucleation. Statistical differences were calculated using the two-sided Mann-Whitney-Wilcoxon test (n = 29 events for APC^Min*/*+^ CTLs and n = 23 events for control CTLs). **(d)** Average engagement parameter of CTLs over the course of the experiment.

## References

[1] K. W. Kinzler and B. Vogelstein, Cell 87, 159 (1996).

[2] C. Luongo, A. R. Moser, S. Gledhill, and W. F. Dove, Cancer Research 54, 5947 (1994).

[3] L. Zhang and J. W. Shay, JNCI: Journal of the National Cancer Institute 109, djw332 (2017).

[4] S. Nelson and I. S. Näthke, Journal of Cell Science 126, 873 (2013).

[5] X. Fang and T. M. Svitkina, European Journal of Cell Biology 101, 151228 (2022).

[6] S. Etienne-Manneville and A. Hall, Nature 421, 753 (2003).

[7] M. Mastrogiovanni, M. Juzans, A. Alcover, and V. Di Bartolo, Frontiers in Cell and Developmental Biology 8 (2020).

[8] A. R. Moser, H. C. Pitot, and W. F. Dove, Science 247, 322 (1990).

[9] L.-K. Su, K. W. Kinzler, B. Vogelstein, A. C. Preisinger, A. R. Moser, C. Luongo, K. A. Gould, and W. F. Dove, Science 256, 668 (1992).

[10] Y. Yamada, K. Hata, Y. Hirose, A. Hara, S. Sugie, T. Kuno, N. Yoshimi, T. Tanaka, and H. Mori, Cancer Research 62, 6367 (2002).

[11] S. Agüera-González, O. T. Burton, E. Vázquez-Chávez, C. Cuche, F. Herit, J. Bouchet, R. Lasserre, I. del Río-Iñiguez, V. Di Bartolo, and A. Alcover, Cell Reports 21, 181 (2017).

[12] M. Juzans, C. Cuche, T. Rose, M. Mastrogiovanni, P. Bochet, V. Di Bartolo, and A. Alcover, ImmunoHorizons 4, 363 (2020).

[13] M. Mastrogiovanni, P. Vargas, T. Rose, C. Cuche, E. Esposito, M. Juzans, H. Laude, A. Schneider, M. Bernard, S. Goyard, C. Renaudat, M.-N. Ungeheuer, J. Delon, A. Alcover, and V. Di Bartolo, Science Advances 8, eabl5942 (2022).

[14] C. Cuche, M. Mastrogiovanni, M. Juzans, H. Laude, M.-N. Ungeheuer, D. Krentzel, M. I. Gariboldi, D. Scott-Algara, M. Madec, S. Goyard, C. Floch, G. Chauveau-Le Friec, P. Lafaye, C. Renaudat, M. Le Bidan, C. Micallef, S. Schmutz, S. Mella, S. Novault, M. Hasan, D. Duffy, V. Di Bartolo, and A. Alcover, Frontiers in Immunology 14 (2023), 10.3389/fimmu.2023.1163466.

[15] S. Budhu, D. A. Schaer, Y. Li, R. Toledo-Crow, K. Panageas, X. Yang, H. Zhong, A. N. Houghton, S. C. Silverstein, T. Merghoub, and J. D. Wolchok, Science Signaling 10, eaak9702 (2017).

[16] B. Weigelin, A. T. den Boer, E. Wagena, K. Broen, H. Dolstra, R. J. de Boer, C. G. Figdor, J. Textor, and P. Friedl, Nature Communications 12, 5217 (2021).

[17] T. E. Schnalzger, M. H. de Groot, C. Zhang, M. H. Mosa, B. E. Michels, J. Röder, T. Darvishi, W. S. Wels, and H. F. Farin, The EMBO Journal 38, e100928 (2019).

[18] J. L. Galeano Niño, S. V. Pageon, S. S. Tay, F. Colakoglu, D. Kempe, J. Hywood, J. K. Mazalo, J. Cremasco, M. A. Govendir, L. F. Dagley, K. Hsu, S. Rizzetto, J. Zieba, G. Rice, V. Prior, G. M. O’Neill, R. J. Williams, D. R. Nisbet, B. Kramer, A. I. Webb, F. Luciani, M. N. Read, and M. Biro, eLife 9, e56554 (2020).

[19] S. Halle, K. A. Keyser, F. R. Stahl, A. Busche, A. Marquardt, X. Zheng, M. Galla, V. Heissmeyer, K. Heller, J. Boelter, K. Wagner, Y. Bischoff, R. Martens, A. Braun, K. Werth, A. Uvarovskii, H. Kempf, M. Meyer-Hermann, R. Arens, M. Kremer, G. Sutter, M. Messerle, and R. Förster, Immunity 44, 233 (2016).

[20] R. Khazen, M. Cazaux, F. Lemaître, B. Corre, Z. Garcia, and P. Bousso, The EMBO Journal 40, e106658 (2021).

[21] G. Ortiz-Muñoz, M. Brown, C. B. Carbone, X. Pechuan-Jorge, V. Rouilly, H. Lindberg, A. T. Ritter, G. Raghupathi, Q. Sun, T. Nicotra, S. R. Mantri, A. Yang, J. Doerr, D. Nagarkar, S. Darmanis, B. Haley, S. Mariathasan, Y. Wang, C. Gomez-Roca, C. E. de Andrea, D. Spigel, T. Wu, L. Delamarre, J. Schöneberg, Z. Modrusan, R. Price, S. J. Turley, I. Mellman, and C. Moussion, Nature 618, 827 (2023).

[22] A. Boussommier-Calleja, R. Li, M. B. Chen, S. C. Wong, and R. D. Kamm, Trends in Cancer 2, 6 (2016).

[23] T. I. Maulana, E. Kromidas, L. Wallstabe, M. Cipriano, M. Alb, C. Zaupa, M. Hudecek, B. Fogal, and P. Loskill, Advanced Drug Delivery Reviews 173, 281 (2021).

[24] S. E. Shelton, H. T. Nguyen, D. A. Barbie, and R. D. Kamm, iScience 24 (2021), 10.1016/j.isci.2020.101985.

[25] M. Campisi, S. E. Shelton, M. Chen, R. D. Kamm, D. A. Barbie, and E. H. Knelson, Cancers 14, 3561 (2022).

[26] S. Sart, G. Ronteix, S. Jain, G. Amselem, and C. N. Baroud, Chemical Reviews 122, 7061 (2022).

[27] R. W. Jenkins, A. R. Aref, P. H. Lizotte, E. Ivanova, S. Stinson, C. W. Zhou, M. Bowden, J. Deng, H. Liu, D. Miao, M. X. He, W. Walker, G. Zhang, T. Tian, C. Cheng, Z. Wei, S. Palakurthi, M. Bittinger, H. Vitzthum, J. W. Kim, A. Merlino, M. Quinn, C. Venkataramani, J. A. Kaplan, A. Portell, P. C. Gokhale, B. Phillips, A. Smart, A. Rotem, R. E. Jones, L. Keogh, M. Anguiano, L. Stapleton, Z. Jia, M. Barzily-Rokni, I. Cañadas, T. C. Thai, M. R. Hammond, R. Vlahos, E. S. Wang, H. Zhang, S. Li, G. J. Hanna, W. Huang, M. P. Hoang, A. Piris, J.-P. Eliane, A. O. Stemmer-Rachamimov, L. Cameron, M.-J. Su, P. Shah, B. Izar, M. Thakuria, N. R. LeBoeuf, G. Rabinowits, V. Gunda, S. Parangi, J. M. Cleary, B. C. Miller, S. Kitajima, R. Thummalapalli, B. Miao, T. U. Barbie, V. Sivathanu, J. Wong, W. G. Richards, R. Bueno, C. H. Yoon, J. Miret, M. Herlyn, L. A. Garraway, E. M. Van Allen,G. J. Freeman, P. T. Kirschmeier, J. H. Lorch, P. A. Ott, F. S. Hodi, K. T. Flaherty, R. D. Kamm, G. M. Boland, K.-K. Wong, D. Dornan, C. P. Paweletz, and D. A. Barbie, Cancer Discovery 8, 196 (2018).

[28] A. W. J. van Renterghem, J. van de Haar, and E. E. Voest, Nature Reviews Clinical Oncology 20, 305 (2023).

[29] M. Nguyen, A. D. Ninno, A. Mencattini, F. Mermet-Meillon, G. Fornabaio, S. S. Evans, M. Cossutta, Y. Khira, W. Han, P. Sirven, F. Pelon, D. D. Giuseppe, F. R. Bertani, A. Gerardino, A. Yamada, S. Descroix, V. Soumelis, F. Mechta-Grigoriou, G. Zalcman, J. Camonis, E. Martinelli, L. Businaro, and M. C. Parrini, Cell Reports 25, 3884 (2018).

[30] J. F. Dekkers, M. Alieva, A. Cleven, F. Keramati, K. L. Wezenaar, E. J. van Vliet, J. Puschhof, P. Brazda, I. Johanna, A. D. Meringa, H. G. Rebel, M.-B. Buchholz, M. Barrera Román, A. L. Zeeman, S. de Blank, D. Fasci, M. H. Geurts, A. M. Cornel, E. Driehuis, R. Millen, T. Straetemans, M. J. T. Nicolasen, T. Aarts-Riemens, H. C. R. Ariese, H. R. Johnson, R. L. van Ineveld, F. Karaiskaki, O. Kopper, Y. E. Bar-Ephraim, K. Kretzschmar, A. M. M. Eggermont, S. Nierkens, E. J. Wehrens, H. G. Stunnenberg, H. Clevers, J. Kuball, Z. Sebestyen, and A. C. Rios, Nature Biotechnology 41, 60 (2022).

[31] G. Ronteix, S. Jain, C. Angely, M. Cazaux, R. Khazen, P. Bousso, and C. N. Baroud, Nature Communications 13, 3111 (2022).

[32] B. Weigelin, E. Bolaños, A. Teijeira, I. Martinez-Forero, S. Labiano, A. Azpilikueta, A. Morales-Kastresana, J. I. Quetglas, E. Wagena, A. R. Sánchez-Paulete, L. Chen, P. Friedl, and I. Melero, Proceedings of the National Academy of Sciences 112, 7551 (2015).

[33] R. F. X. Tomasi, S. Sart, T. Champetier, and C. N. Baroud, Cell Reports 31, 107670 (2020).

[34] P. Abbyad, R. Dangla, A. Alexandrou, and C. N. Baroud, Lab Chip 11, 813 (2011).

[35] S. Sart, R. F.-X. Tomasi, G. Amselem, and C. N. Baroud, Nature Communications 8, 469 (2017).

[36] S. Sart, R. F.-X. Tomasi, A. Barizien, G. Amselem, A. Cumano, and C. N. Baroud, Science Advances 6, eaaw7853 (2020).

[37] T. Ohira, Y. Ohe, Y. Heike, E. R. Podack, K. J. Olsen, K. Nishio, M. Nishio, Y. Miyahara, Y. Funayama, H. Ogasawara, H. Arioka, H. Kunikane, M. Fukuda, H. Kato, and N. Saijo, Journal of Cancer Research and Clinical Oncology 120, 631 (1994).

[38] B. Breart, F. Lemaître, S. Celli, and P. Bousso, The Journal of Clinical Investigation 118, 1390 (2008).

[39] K. Lei, A. Kurum, M. Kaynak, L. Bonati, Y. Han, V. Cencen, M. Gao, Y.-Q. Xie, Y. Guo, M. T. M. Hannebelle, Y. Wu, G. Zhou, M. Guo, G. E. Fantner, M. S. Sakar, and L. Tang, Nature Biomedical Engineering 5, 1411 (2021).

[40] M. Desouza, P. W. Gunning, and J. R. Stehn, BioArchitecture 2, 75 (2012).

[41] J. P. Doncel, P. de la Cruz Ojeda, M. OropesaÁvila, M. V. Paz, I. D. Lavera, M. D. L. Mata, M. Álvarez Córdoba, R. L. Hidalgo, J. M. S. Rivero, D. Cotán, and J. A. Sánchez-Alcázar, in Cytoskeleton, edited by J. C. Jimenez-Lopez (IntechOpen, Rijeka, 2017) Chap. 8.

[42] T. H. Harris, E. J. Banigan, D. A. Christian, C. Konradt, E. D. Tait Wojno, K. Norose, E. H. Wilson, B. John, W. Weninger, A. D. Luster, A. J. Liu, and C. A. Hunter, Nature 486, 545 (2012).

[43] E. R. Jerison and S. R. Quake, eLife 9, e53933 (2020).

[44] M. G. Overstreet, A. Gaylo, B. R. Angermann, A. Hughson, Y.-M. Hyun, K. Lambert, M. Acharya, A. C. Billroth-MacLurg, A. F. Rosenberg, D. J. Topham, H. Yagita, M. Kim, A. Lacy-Hulbert, M. Meier-Schellersheim, and D. J. Fowell, Nature Immunology 14, 949 (2013).

[45] M. P. Matheu, C. Beeton, A. Garcia, V. Chi, S. Rangaraju, O. Safrina, K. Monaghan, M. I. Uemura, D. Li, S. Pal, L. M. De La Maza, E. Monuki, A. Flügel, M. W. Pennington, I. Parker, K. G. Chandy, and M. D. Cahalan, Immunity 29, 602 (2008).

[46] E. H. Wilson, T. H. Harris, P. Mrass, B. John, E. D. Tait, G. F. Wu, M. Pepper, E. J. Wherry, F. Dzierzinski, D. Roos, P. G. Haydon, T. M. Laufer, W. Weninger, and C. A. Hunter, Immunity 30, 300 (2009).

[47] H. Salmon, K. Franciszkiewicz, D. Damotte, M.-C. Dieu-Nosjean, P. Validire, A. Trautmann, F. Mami-Chouaib, and E. Donnadieu, The Journal of Clinical Investigation 122, 899 (2012).

[48] J. B. Beltman, A. F. M. Marée, and R. J. de Boer, Nature Reviews Immunology 9, 789 (2009).

[49] R. Thibaut, P. Bost, I. Milo, M. Cazaux, F. Lemaître, Z. Garcia, I. Amit, B. Breart, C. Cornuot, B. Schwikowski, and P. Bousso, Nature Cancer 1, 302 (2020).

[50] T. Song, Y. Choi, J.-H. Jeon, and Y.-K. Cho, Frontiers in Immunology 14 (2023).

[51] S. Agüera-Gonzalez, J. Bouchet, and A. Alcover, in eLS (John Wiley & Sons, Ltd, 2015).

[52] M. de la Roche, Y. Asano, and G. M. Griffiths, Nature Reviews Immunology 16, 421 (2016).

[53] M. Barry and R. C. Bleackley, Nature Reviews Immunology 2, 401 (2002).

[54] M. L. Dustin, S. K. Bromley, Z. Kan, D. A. Peterson, and E. R. Unanue, Proceedings of the National Academy of Sciences 94, 3909 (1997).

[55] B. Weigelin and P. Friedl, Trends in Cancer 8, 980 (2022).

[56] M. Scabini, F. Stellari, P. Cappella, S. Rizzitano, G. Texido, and E. Pesenti, Apoptosis 16, 198 (2011).

[57] H. D. Moreau, F. Lemaître, E. Terriac, G. Azar, M. Piel, A.-M. Lennon-Dumenil, and P. Bousso, Immunity 37, 351 (2012).

[58] G. Amselem, S. Sart, and C. N. Baroud, in Methods in Cell Biology, Microfluidics in Cell Biology Part C: Microfluidics for Cellular and Subcellular Analysis, Vol. 148, edited by D. A. Fletcher, J. Doh, and M. Piel (Academic Press, 2018) pp. 177–199.

[59] D. Ershov, M.-S. Phan, J. W. Pylvänäinen, S. U. Rigaud, L. Le Blanc, A. Charles-Orszag, J. R. W. Conway, R. F. Laine, N. H. Roy, D. Bonazzi, G. Duménil, G. Jacquemet, and J.-Y. Tinevez, Nature Methods 19, 829 (2022).

